# On the potential for GWAS with phenotypic population means and allele-frequency data (popGWAS)

**DOI:** 10.1101/2024.06.12.598621

**Authors:** Markus Pfenninger

## Abstract

It is vital to understand the genomic basis of differences in ecologically important traits if we are to understand the impact of global change on biodiversity and enhance our ability for targeted intervention. This study explores the potential of a novel genome-wide association study (GWAS) approach for identifying loci underlying quantitative polygenic traits in natural populations, based on phenotypic population means and genome-wide allele frequency data as obtained e.g. by PoolSeq approaches. Extensive population genetic forward simulations demonstrate that the approach is generally effective for oligogenic and moderately polygenic traits and relatively insensitive to low heritability. However, applicability is limited for highly polygenic architectures and pronounced population structure. The required sample size is moderate with very good results being obtained already for a few dozen populations scored. When combined with machine learning for feature selection, the method performs very well in predicting population means. The data efficiency of the method, particularly when using pooled sequencing and bulk phenotyping, makes GWAS studies more accessible for research in biodiversity genomics. Moreover, in a direct comparison to individual based GWAS, the proposed method performed constistently better with regard to the number of true positive loci identified and prediction accuracy. Overall, this study highlights the promise of popGWAS for dissecting the genetic basis of complex traits in natural populations.

## Introduction

A major goal as well as a major challenge in evolutionary biology is to understand how genes influence traits, i.e. the genotype-phenotype link, (Brandes et al., 2022; Uffelmann et al., 2021). The difficulties in achieving this goal are primarily due to the fact that the heritable variation of many, if not most, relevant phenotypes is determined by small contributions from many genetic loci (Sella & Barton, 2019). Such complex traits are usually influenced by a few dozen genes that are mechanistically directly involved in their expression, but often also by numerous, if not almost all, other genes if only by the use of common resources and machinery (omnigenic theory) (Boyle et al., 2017) as well as the environment in the widest sense.

Genome-wide association studies (GWAS) are commonly used to link complex phenotypic traits to their genomic basis (Brandes et al., 2022; Visscher et al., 2012). However, given the complexity of causal mechanisms and the small effects of individual loci, often only a small fraction of the genetic variation underlying phenotypic variance is identified, despite the considerable logistic effort in terms of the number of phenotyped and genotyped individuals (Brandes et al., 2022; Visscher et al., 2017). As a result, accurate predictions of phenotypes from genomic data are still quite limited and there is currently no other strategy than to keep increasing sample sizes (Brandes et al., 2022). This is a problem in the medical sciences (Shendure et al., 2019), but the greater challenge for science and society probably lies in addressing the global biodiversity crisis. It would be highly desirable to have affordable methods to accurately understand the genomic basis of relevant traits and predict (non-model) species responses to all aspects of global change (Bernatchez et al., 2023; Waldvogel et al., 2020).

GWAS with wild populations has been advocated for some time (Santure & Garant, 2018). However, despite recent progress in high throughput, automated phenotyping (Dunker et al., 2022; Tills et al., 2023; Xie & Yang, 2020), the advances of biodiversity genomics in obtaining high quality reference genomes for almost every species (Exposito-Alonso et al., 2020; Formenti et al., 2022) and the possibility to gain cost-effective genome-wide population data (Czech, Peng, Spence, Lang, Bellagio, Hildebrandt, Fritschi, Schwab, Rowan, & Weigel, 2022; Schlötterer et al., 2014), relatively few empirical studies are currently available (e.g. Gauzere et al., 2023; Ithnin et al., 2021; James et al., 2022). This gap between the possibilities and actual practical application in biodiversity conservation (Heuertz et al., 2023; Hogg, 2023) is probably as much due to the still existing logistic and financial challenges as to a lack of data- and resource-efficient methods.

Here, I explore the potential of a new GWAS approach using phenotypic population means and genome-wide allele-frequency data. The rationale behind the approach is straightforward. If a quantitative polygenic trait has an additive genetic component, an individual’s phenotypic trait value should at least roughly correlate with the number of trait-increasing alleles at the underlying loci (Uffelmann et al., 2021). Consequently, it was theoretically expected (Orr, 1998; Pritchard & Di Rienzo, 2010) and empirically shown (Turchin et al., 2012) that trait-increasing alleles will tend to have greater frequencies in the population with higher mean trait values, compared to the population with a lower trait mean. When examining populations with a range of different phenotypic trait means, we may therefore expect that the allele frequencies at the trait-affecting loci show a linear or at least steady relation with the observed trait means (Barton, 1999). I hypothesise here that this predicted relation can be exploited to distinguish potentially causal loci (and the linked variation) from loci not associated with the focal trait. In case of a successful evaluation, the major advantages of the proposed approach would be the reduced sequencing effort by the possibility to use pooled population samples (PoolSeq) and the opportunity to use bulk phenotyping (e.g. by satellite imaging, flow-cytometry, etc.) on traits for which individual phenotyping is difficult or tedious.

The most important assumption for the approach is obviously that observed population differences in the focal trait means have at least partially a genetic basis. Since the environment has usually an effect on the phenotype (Sella & Barton, 2019), total phenotypic variance should be adjusted for known fixed environmental effects, because this increases the fraction of variance due to genetic factors (Visscher et al. 2008). Predicting additive genetic values with even higher accuracy can be achieved by taking into account GxE interactions through repeated phenotypic measurements of the same individuals under different environmental conditions, e.g. by time series (Visscher et al. 2008). I assumed therefore that environmental influence on the phenotypic trait variance among populations has been statistically removed as much as possible (Harpak & Przeworski, 2021). Similarly important is the assumption that the genetic variance of the focal quantitative trait can be adequately described by an additive model. Both empirical and theoretical evidence suggests that this is indeed the case for most complex traits (Hill et al., 2008). Even though epistatic interactions are wide spread (Mackay, 2014), Sella and Barton (Sella & Barton, 2019) argue that the marginal allelic effects on quantitative traits are well approximated by a simple additive model.

The aims of this study were i) to understand whether and under which circumstances the hypothesised pattern of a linear relation between the population allele frequencies at causal loci and the phenotypic population means of the respective trait emerges, ii) to evaluate the influence of population genetic parameters of typical natural systems and the experimental design on the likelihood of identifying causal loci underlying an additive quantitative trait, in particular to elucidate the limits of the approach with regard to genetic architecture and population structure, iii) to compare proposed method with the performance of an individual-based GWAS, iv) to explore the possibilities for statistical genomic prediction of phenotypic population means from the allele frequencies at identified loci, and v) to evaluate the statistical power of the method for a realistic range of effective genome sizes. I used individual-based population genomic forward simulations and machine learning approaches (minimum entropy feature selection) for prediction and utilised an information theory-based framework for evaluation of the proposed method.

## Material and methods

### Rationale of the method

#### Expectation of a positive correlation between quantitative trait loci allele frequencies and phenotypic population means

Consider a biallelic, codominant system for the additively heritable component of a quantitative trait with *n* loci contributing to the trait. In this system, all loci contribute equally to the phenotypic trait, with one allele per locus making a greater contribution than the other. The phenotypic trait value *x* of an individual can then be determined by simply adding up the number of trait-increasing alleles (*g* with values of 0, 1 or 2) over all *n* quantitative trait loci (QTL) and multiplying this sum with a scaling constant *k*:

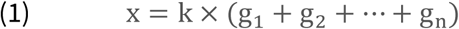

When adding more individuals, the phenotypic population trait mean is defined as the mean of the row sums:

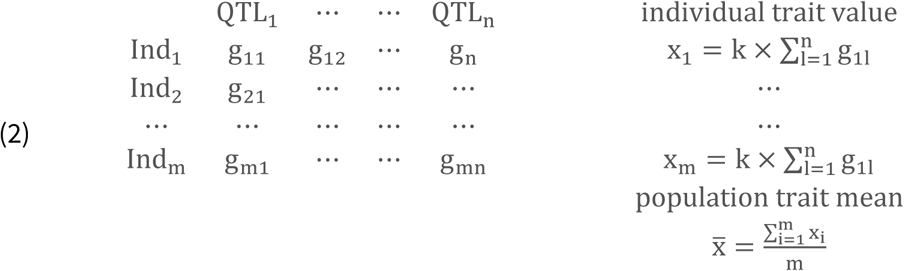

The columns of this matrix can be used to calculate the population allele frequency (AF) of the trait increasing allele for each QTL.

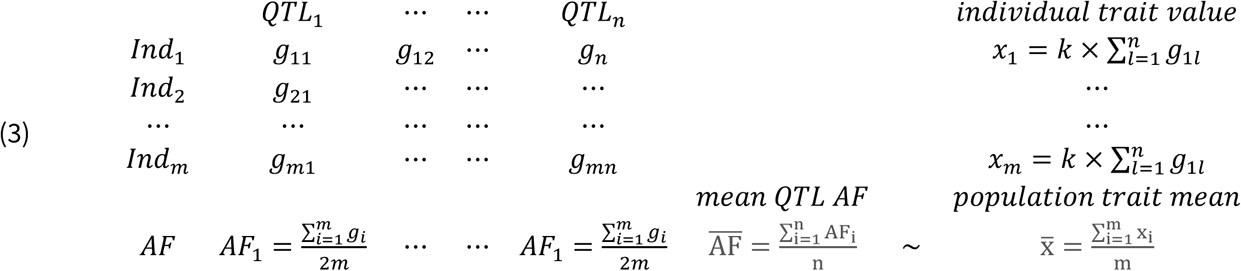

The mean population allele frequency at the QTL loci is thus directly proportional to the phenotypic population trait mean. This relationship remains unchanged even if the individual locus contributions are not identical, with some loci contributing more or less to the phenotypic trait value. In this case, a scaling vector is required to weigh the individual locus contributions to individual trait values, and those of the AFs to the population trait mean. Since the AFs are by definition bounded by zero and one, the population trait mean is minimal when the allele frequencies of the trait-increasing allele at all QTL are zero and maximal when all QTL AFs are one. This proportionality links the individual genotypes and the AFs at the QTL linearly with the population trait mean.

If we extend this to a set of populations and order them with decreasing phenotypic population means, we can be sure that the mean QTL AFs of the populations will also be ordered in decreasing sequence:

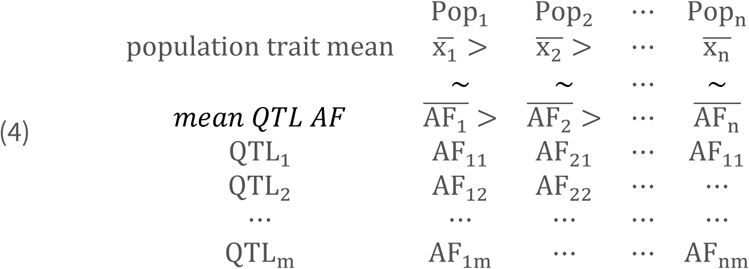

The answer to whether the allele frequencies in every row i.e. at every contributing locus can be used to predict the population trait mean depends on whether the expected covariance between these two vectors is positive:

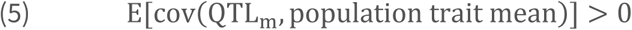

where

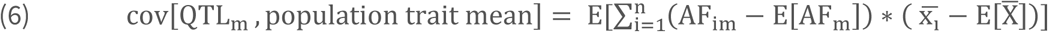

with X̅ representing the grand mean over all populations. As all elements in the QTL matrix are positive, they inherently tend to contribute positively to their column means. Therefore, AF larger than the overall AF mean at his locus tend to be on the left side of the population closest to the overall phenotypic mean in the ordered matrix above. Conversely, AF smaller than the locus AF mean are rather on the right. This leads intuitively to a positive expected covariance between each row and the column mean, in particular if the number of populations becomes large. Conversely, the AF at (unlinked) loci not contributing to the phenotypic population trait mean have an expectation of zero.

### Testing the expectations of a detectable linear relation between allele frequencies and phenotypic population means

I tested these general expectations and the effect of different scaling vectors for the effect size distribution of QTL with a first set of simulations. I generated a matrix of size *n x m* populated with random AF between zero and 1. To avoid stochastic effects due to sample size, the number of populations *n* was fixed at 10,000. The number of QTL *m* was varied from oligogenic to highly polygenic (10, 20, 50, 100, 200, 500, 1000, 2000, 5000). Three different distributions of loci effects were tested, i) a flat distribution with all loci contributing equally, ii) a mildly decreasing exponential function and iii) a steeply decreasing exponential function with few loci contributing much and many very little (Supplemental Figure 1).

Each of the *m* columns was used to calculate the phenotypic population mean of the respective population by adding up the AF multiplied with the respective locus weight. The resulting *n* phenotypic population means were then correlated to the *n* AF of each of the *m* loci and the resulting *m* Pearson correlation coefficients (i.e. the standardized covariance) recorded. From these, mean and standard deviation were calculated and tested, whether they conform to a normal distribution (scipy.stats.normaltest). Furthermore, a second matrix of identical size was populated with random AF, and the correlation of these non-contributing loci to the population means derived from the QTL matrix was computed. The simulations were repeated 10 times in every possible parameter combination and the results averaged (Supplemental Script 1).

### Individual based Wright-Fisher forward model used for simulation

A Wright-Fisher individual forward genetic simulation model was used to investigate the potential of a genome-wide association study based on the means of a population trait and population allele frequency data. As this model formed the basis for all further simulations, its general structure is first described.

In the simulation, all loci were assumed to be unlinked, thus representing haplotypes in LD rather than single SNPs. (Visscher et al., 2017). For each simulation run, the initial allele frequencies for all loci in the total population were randomly drawn from a beta function with parameters α = β = 0.5 in a range 0.1 to 0.9. To generate a hermaphroditic and diploid individual, two alleles were randomly drawn with a probability based on their frequency at the respective locus, and the resulting genotype at this locus was recorded. This process was repeated for all loci. As a result, each individual was represented by a vector of biallelic genotypes (*AA* = 0, *Aa*;*aA* = 1, *aa* = 2). To model a quantitative, fully additive trait, a variable number of loci were assigned as quantitative trait loci (QTL). In addition, a much larger number of neutral loci was modelled.

#### Genetic architecture of the quantitative trait

The continuous trait value was measured in arbitrary units. The allele (*A*) at each QTL had no effect on the individual’s trait value, resulting in a completely homozygous *A* individual at the QTL having a phenotypic trait value of 0. The alternative allele (*a*) added a locus-specific value to the trait. Two distributional extremes have been considered for the allelic effects on the trait value: i) a uniform distribution where each locus contributes 0.5 units to the trait. An individual that is completely homozygous for the alternative allele *a* therefore had a trait value equal to the number of QTL, ii) an exponential distribution with few loci having large effects and many having very small effects, scaled such that the maximum possible trait value was also equal to the number of QTL (see Supplementary Figure 2A). To model the effect of phenotyping errors, unaccounted environmental influence, and/or the unspecific contribution of the genomic background to the trait, a random value drawn from a Gaussian distribution with a mean of zero and selectable standard deviation between 0.1 and 3 was added to the genetically determined phenotype value of each individual. The phenotypic value of each individual’s trait was determined by summing the allelic effects of all genotypes at all QTL loci plus the random value and the result recorded.

It was assumed that all trait variation was based on standing genetic variation. Mutations were not considered because new or low frequency mutations do not substantially influence phenotypic population means and their AFs. In addition, mutations are rare and mutations affecting a particular functional trait even rarer. Therefore, restricting the simulations to standing genetic variation seemed reasonable.

#### Reproduction and selection

Subpopulations in each run were created from the same initially drawn random allele frequency array, mimicking a common descent. Due to sampling variance, the realised allele frequencies and thus the mean subpopulation trait value differed from the initial frequencies of the total (ancestral) population. A subpopulation always comprised 500 adult individuals.

For reproduction, two random individuals were chosen with replacement from the adult population. The genotype of an offspring individual at a locus was determined by randomly choosing one of the two alleles from each designated parent at this locus. Each parent fostered n_juv offspring; therefore, 2 x n_juv were produced in each mating. After reproduction, the parental generation was discarded to prevent overlapping generations. Each generation had N/2 matings, resulting in an offspring population of N*n_juv individuals. This assured that the genotypes in each subpopulation were in Hardy-Weinberg equilibrium after the first generation.

Because the offspring population was much larger than the size of the adult population, it was necessary to reduce it. This was achieved by a combination of ‘hard’ natural selection and random mortality. An individual’s survival to the adult stage was determined by the absolute deviation of its phenotypic trait value from a pre-specified selective trait optimum for the respective subpopulation. This selective trait optimum for a subpopulation was determined by adding a random value taken from a Gaussian distribution with a mean of zero and a standard deviation of 2.5 to the initial population mean. An individual’s survival probability was determined by an exponential decline function with strength *s* (the exponent of the function, see Supplementary Figure 2B). Individuals were randomly selected one by one from the offspring population, the distance of their phenotype to the selective optimum calculated and their survival probability calculated. A respectively biased coin was then tossed to determine their fate. This process was repeated until the adult population size was reached, and any remaining offspring individuals were indiscriminately discarded. If the phenotypic mean of the subpopulation was close to or at the selective optimum (see below), this process resulted in stabilising selection. If the population was away from the optimum, rapid directed selection towards the optimum was observed, depending on the strength of selection.

#### Population structure

For the assessment of the effect of population structure, each subpopulation received in every generation a certain number of migrants randomly chosen from all other subpopulations. Both drift and selection towards different trait optima led to variation in population trait means among the subpopulations.

Based on the model described above, I considered scenarios were subpopulations with quantitative phenotypic population differences in mean for the trait in question were screened from a larger total population. Although the population trait mean differences in the simulation of this scenario were created by drift and local adaptation, any other source of heritable phenotypic population differentiation, such as maladaptation, introgression, or e.g. in the case of managed species, human choice, may also be the non-exclusive reasons for differentiation in population means. The range of phenotypic variation among the subpopulations was not predetermined, but an emergend feature of the simulation parameters.

After evolving the subpopulations for the desired number of generations, phenotypic trait means, and genome-wide allele frequencies were recorded. While the phenotypic means for each subpopulation was calculated over all individuals, the allele frequencies were estimated in a PoolSeq (Kofler et al., 2011) like fashion from subsamples of 50 individuals. The range of phenotypic trait means of the population sample was recorded. Trait heritability was determined in the last generation by regressing the phenotypic values of the offspring against the mean of their respective parents (Lynch & Walsh, 1998). As measure for population subdivision due to drift, F_ST_ among all subpopulations was calculated from the variance of the true allele frequencies (Wright, 1949).

### Simulated scenarios

#### Influence of natural system and experimental design factors

In a first set of simulations, I explored the influence of factors inherent to the natural system and the experimental design on population GWAS performance (Table 1). As factors of the natural system, I assumed characteristics that are beyond control of the researcher, such as heritability of the trait and its genetic architecture (number of QTL, distribution of allele trait contribution). While the degree of population differentiation and range of phenotypic differentiation are also inherent to the organism studied, the choice of samples may allow a certain control over these parameters. The number of subpopulations screened is clearly a study design decision (Table 1).

**Table 1.** Simulation parameters, their abbreviations, values used in simulations, their biological meaning and whether the parameter is a feature of the natural system under scrutiny or under the control of the researcher.

| Parameter | Abbreviation | Values in the simulation | Biological meaning | Degree of knowledge in natural systems/under the control of study design |
| --- | --- | --- | --- | --- |
| Number of subpopulations scored for phenotypic population means and genome wide allele frequencies | n_pop | 12, 24 36, 48, 60 | - | Full control |
| Number of quantitative trait loci contributing to the focal trait | n_qtl | 30, 50, 70, 110, 500 | Genetic architecture of the trait | *A priori* unknown |
| Distribution of allelic effects on the focal trait | allelic_contr | Flat, exponential | Genetic architecture of the trait | *A priori* unknown |
| Standard deviation of random phenotypic variation added to individuals | env-var | 0.1, 1, 2, 3 | Heritability of the trait | *A priori* unknown |
| Number of migrants received in each subpopulation per generation | mig | 0, 5, 10, 50, 100 | Population structure | Partial control |

A genetic trait architecture of 30, 50, 70, 110 and 500 loci, flat and exponential allelic effect distributions, as well as environmental variance coefficients of 0.1, 1, 2 and 3 were applied (Table 1). Selection strength was fixed at 0.5 (Supplemental Figure 2B). The number of immigrants reaching and reproducing in each subpopulation in each generation was varied between 5, 10, 50 and 100 individuals. Simulations were run for 30 generations among 12, 24, 36, 48 and 60 subpopulations of 500 individuals each (Table 1). For this set of simulations, 1000 neutral loci and a fixed outlier threshold (upper 5% quantile, either 21 or 22 outlier loci, respectively) were applied. Each possible parameter combination was run in five replicates, resulting in 4000 simulation runs.

The effect of each parameter on PPV was assessed with ANOVA over all simulations, grouped after the respective parameter classes. The relative influence of the number of populations, QTL loci, distribution of allelic contributions, trait heritability, phenotypic range and population subdivision on the proportion of True Positive Loci (TPL) among the outlier loci was determined with a General Linearized Model (GLM).

#### Hierarchical population structure

The effect of a hierarchical population structure on popGWAS performance was assessed with a restricted parameter set (Table 2). Two, three and four regions were simulated, each consisting of 12 subpopulations. While there was gene-flow among the populations within the regions as described above, no gene-flow among regions was simulated. The initial degree of differentiation among regions was determined by first drawing one set of random (beta distributed) allele frequencies as described above. To start each independent region with slightly, but not completely different allele frequencies, a random variate from a normal distribution with mean 0 and either 0.05, 0.1 or 0.2 as standard deviation was then added to each of the initially drawn allele frequencies (Table 2). The simulations were then run as described above. The hierarchical population structure was not explicitely taken into account in statistical analysis but performed as described above.

**Table 2.** Simulation parameters for hierarchical population structure, their abbreviations, values used in simulations, their biological meaning and whether the parameter is a feature of the natural system under scrutiny or under the control of the researcher.

| Parameter | Abbreviation | Values in the simulation | Biological meaning | Degree of knowledge in natural systems/under the control of study design |
| --- | --- | --- | --- | --- |
| Number of regions with 12 subpopulations each | n_reg | 2,3,4 | - | Partial control |
| Number of quantitative trait loci contributing to the focal trait | n_qtl | 30, 70, 110 | Genetic architecture of the trait | *A priori* unknown |
| Distribution of allelic effects on the focal trait | allelic_contr | Flat, exponential | Genetic architecture of the trait | *A priori* unknown |
| Standard deviation of random phenotypic variation added to individuals | env_var | 2 | Heritability of the trait | *A priori* unknown |
| Number of migrants received in each subpopulation per generation | mig | 5, 50 | Gene-flow | Partial control |

#### Population based GWAS (popGWAS)

Assuming a linear relation between the phenotypic (sub)population means and the population allele frequencies of the causal loci, I calculated an ordinary linear regression between these two variables for all loci in the genome for each simulation. I used the resulting

-log_10_ p value as measure of regression fit and effect size. I recorded the number of true positive loci (TPL) among the loci beyond a predefined outlier threshold (2% quantile). As GWAS performance measures, the true positive rate (TPR = recall, sensitivity, discovered proportion of all QTL), positive predictive value (PPV = precision, proportion of TPL among outliers considered) and false discovery rate (1 – PPV = FDR = proportion of false positive loci among outliers considered, type I error) were calculated. This was performed for all simulated scenarios.

#### Individual GWAS (iGWAS)

In addition, a traditional linear GWAS on a random sample of individuals from all subpopulations was performed for all simulation scenarios (termed iGWAS hereafter). The number of individuals was chosen such that the PoolSeq sequencing effort for the same simulation was matched, assuming that a 15X mean coverage is appropriate for accurate genotyping. A linear regression of the individuals’ genotypes at each locus as predictors for the phenotypes of each individual in the sample was calculated. To assure comparability with the popGWAS approach and keep computation times feasible, no correction method was applied. The resulting -log10p values were treated in the same way as described for popGWAS above.

### Genomic prediction and validation

The loci identified by popGWAS were used to devise a statistical genomic prediction model to obtain a score that uses observed allele frequencies at the identified loci to predict the mean population phenotype of unmeasured populations. To remove remaining uninformative or redundant loci, I applied feature selection, which is particularly suitable for bioinformatic data sets that contain many features but comparatively few data points. The minimum entropy feature selection (MEFS) technique uses mutual information to measure the dependence between each feature and the target variable. For a given number of features (k), the data set of the allele frequencies at selected outlier loci and the respective phenotypic population means was repeatedly randomly divided in training (80%) and test set (20%), a multiple regression model fitted and the r²-fit of the test sets to the predicted phenotypes recorded. The best model for the current k was recorded and the process repeated for all k in a range between 2 and the number of selected loci – 1. Finally, the best model (i.e. highest r²) among all k was chosen as best prediction model. MEFS was implemented with the Python module scikit-learn 1.3.2 (Pedregosa et al., 2011)

The performance of the selected best prediction model for each run was tested with independent data. Ten additional subpopulations were created and evolved under the same parameters as part of the metapopulation of the initial set of populations and their mean population phenotypes calculated as described above. Then the allele frequencies at the predictive loci as identified by the best prediction model were extracted and phenotypic prediction scores according to the best prediction model calculated. The performance of the statistical genomic prediction was then evaluated by calculating the Pearson correlation coefficient *r* between the observed mean population phenotypes and the phenotypic prediction scores for the ten validation populations (Supplemental Script 2).

Genomic prediction and validation were performed in the same manner for iGWAS. Here it was the goal to predict the individual phenotypes of a sample not used to train the model with the genotypes of the loci identified by iGWAS as predictors.

### Method performance with realistic genome sizes

Whether and which proportion of TPL, i.e. causal loci can be expected to be reliably identified with the proposed method depends crucially on the total number of loci screened as this number determines the length and size of the distributional tail of random associations of neutral loci with the mean population phenotypes. The number of effectively independently evolving loci in a population depends on genome size, effective population size (including all factors that affect it locally and globally) and recombination rate (Chakraborty, 1981; Taylor & Higgs, 2000). There are hardly any empirical estimates in the literature, but dividing typical genome sizes by typical mean genome-wide LD ranges suggested that a few tens of thousands to a few hundreds of thousands of independent loci per genome is a realistic range for a large number of taxa (see Supplemental Table 1). I have therefore considered 1,000, 5,000, 10,000, 30,000, 50,000 and 100,000 independent neutral loci for samples of 12, 24, 36, 48 and 60 populations with a restricted set of parameters (number of QTL and allelic contribution). As the true number of QTL underlying a trait is rarely *a priori* known, I considered 10, 30, 50, 70 and 110 QTL loci in this analysis. I therefore recorded the number of TPL found in sets of loci with the absolutely highest 10, 30, 50 and 100 -log_10_p values, as well as outlier proportions of 0.0001, 0.001, 0.01, 0.02, 0.05 and 0.1 of the total number of loci in the respective simulation. As above, all simulations were run in all possible parameter combinations with five replicates each (Supplemental Script 3).

I analysed the performance of the method in an Area Under the Curve – Receiver Operator Curve (AUC-ROC) and – Precision, Recall (AUC-PR) framework as suggested by Lotterhos et al. (Lotterhos et al., 2022). For each combination of effective genome size and number of population scored, mean TPR, PPV and FDR were calculated over all replicates and parameter combinations for the respective set of simulations. The maximum F1 score (Rijsbergen, 1979), the harmonic mean of the precision and recall representing both in one metric, was used in addition to identify the optimal number of outliers to select.

All simulations were implemented in Python 3.11.7 (Van Rossum & Drake, 2009) and run under pypy 3.10 (Team, 2019), the respective scripts can be found in the Supplementary Material (Scripts 1-3). General statistical tests were performed with R (R Core Team, 2013).

## Results

### Allele frequencies at QTL loci co-vary positively with the population trait mean

The mean correlation coefficient between all QTL and the respective phenotypic population means was positive in every parameter combination and in every single simulation (Table 3).

**Table 3.**
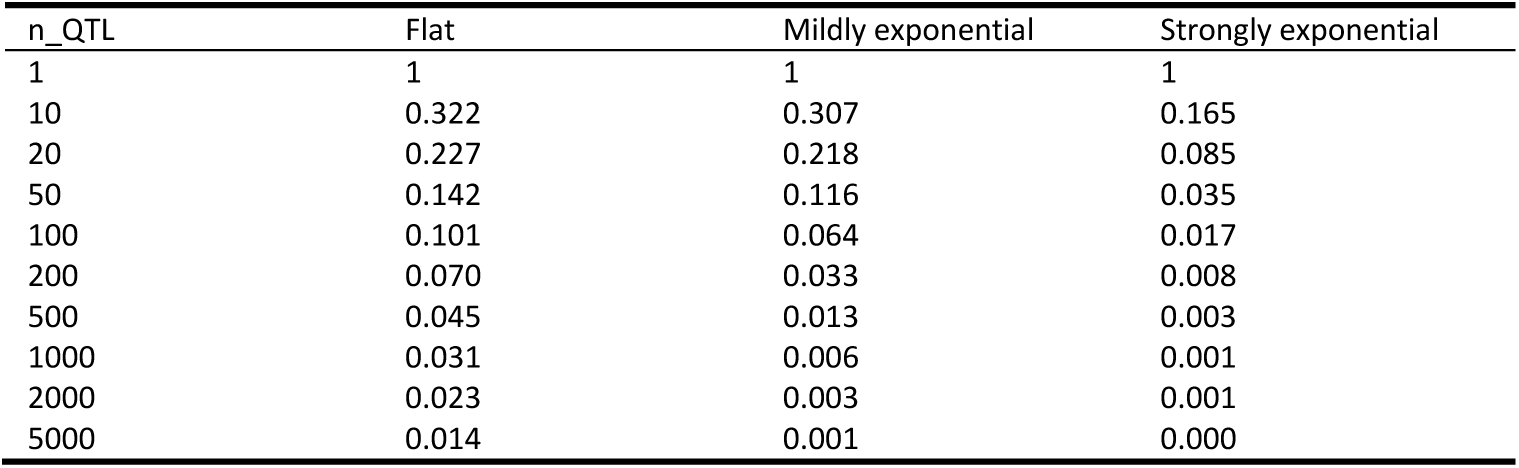
Expected mean Pearson’s correlation coefficients between QTL AF and phenotypic population means for three different locus contribution distributions and varying number of QTL.

The expected mean correlation coefficient decreased with increasing number of contributing QTL (Figure 1). This decay was best described by a negative exponential function of the form *number of QTL^-1/x^* with x ranging from 1.26 in case of the strongly unbalanced locus contributions to 2 for the flat distribution.

**Figure 1.**
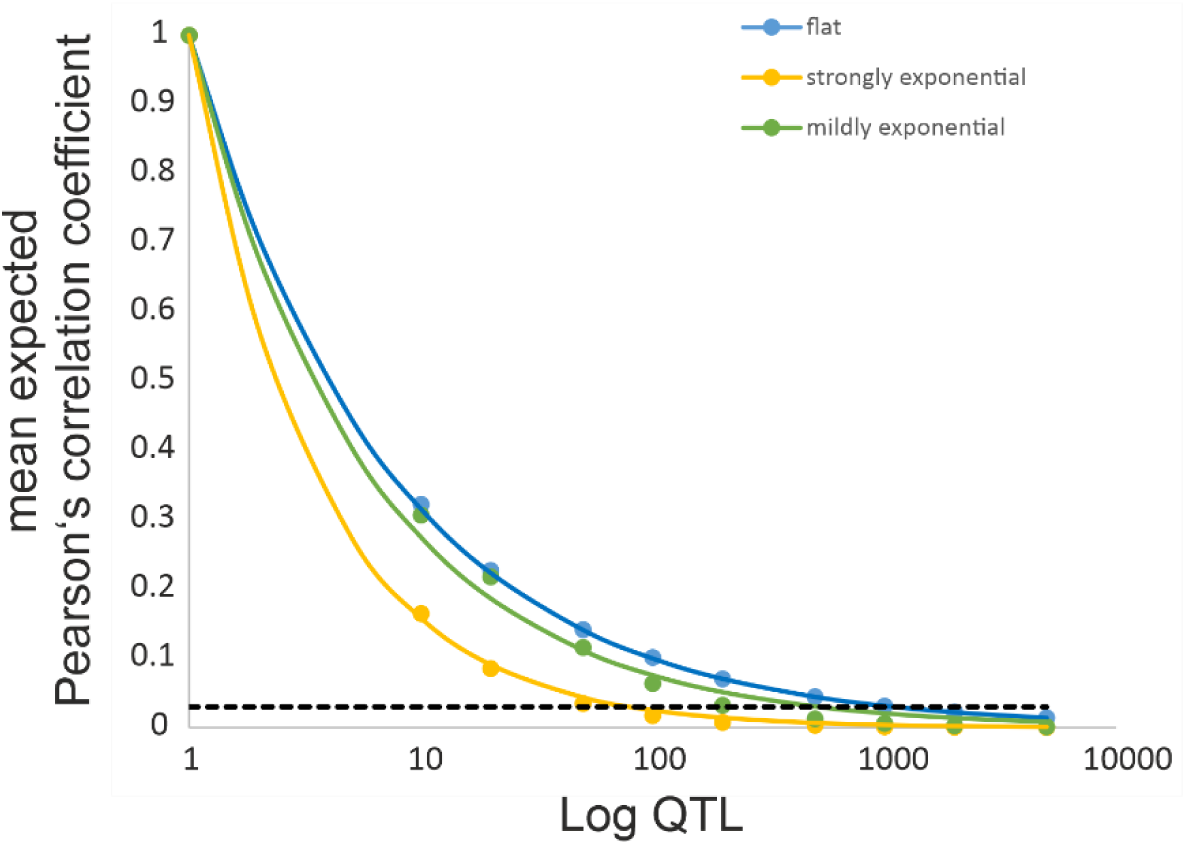
Plot of the expected correlation coefficient between individual QTL loci and the population trait mean in dependence of the number of QTL loci. The dashed black line gives the upper 95% confidence interval for non-contributing, neutral loci.

The distribution of the correlation coefficients did not deviate from a normal distribution for the flat locus contribution distribution, while it did for all other parameters. The correlation coefficients for the non-contributing loci had an expectation of zero and a mean standard deviation of 0.01, regardless of the number of loci.

### Influence of simulation parameters on parameters of the simulated populations

In an initial set of simulations, the effect of the number of generations simulated and the number of migrants on the degree of adaptation and population structure was simulated. It turned out that selection/drift equilibrium was reached after less than 10 generations. As expected, more gene-flow led to less perfect local adaptation (Supplementary Figure 3A). The time to reach the equilibrium F_ST_ among subpopulations strongly depended on the level of gene-flow. While it was swiftly reached within 50 generations for 20 or more migrants per generation, it would have taken much longer for less gene-flow (Supplementary Figure 3B). However, popGWAS performance was governed by the F_ST_ at the time of analysis and not by selection-migration-drift equilibrium (Supplemental Figure 3C). To keep computation times for the individual based model within reasonable bounds, all following simulations were run for 30 generations.

In the second set of simulations with 1000 neutral loci and an outlier threshold of the most extreme 2% -log_10_p values, the permutation of all parameters with five replicates yielded 3884 completed independent simulation runs. The 116 missing simulations to the expected 4000 runs were due to one or more subpopulations going extinct during the simulation.

The number of migrants strongly influenced the population structure (r² = 0.990). F_ST_ estimates decreased exponentially(Supplemental Figure 4A). Heritability of the trait depended strongly on the environmental variation parameter (r² = 0.788, Supplemental Figure 4B). It decreased on average by 0.18 per unit standard deviation, with variations of up to 0.05 even among runs with identical parameters. Trait heritability estimates ranged from 0.21 to 1.02.

### Factors influencing the proportion of detected true positive loci among outliers

Over all simulations in the first set of scenarios, on average about 12.1 (mean proportion 0.52) true positive loci (TPL) were among the highest 2% outliers. The TPL values ranged between none (0) and 23 (1.0); the 25 percentile was 6 (0.28), the 75 percentile 19 (0.81). This exceeded in >90% of cases random expectations, when excluding the highly polygenic case (n_qtl = 500), this proportion rose to more than 93%. No p-value inflation was observed; the mean slope of qq-plots was 1.27 (s.d. 0.23), the distribution ranged between 1 and 1.99 (Supplemental Figure 4A). The slope of the qq-plots was mainly driven by the number of TPL detected (r = 0.65, p = 0). Supplemental Figure 5 illustrates exemplarily the results of three individual simulation runs with different parameter sets.

The environmental variability parameter had no significant effect on PPV (F = 0.13, p = 0.939, Figure 2A). The number of QTL showed a systematic effect on mean PPV (ANOVA F = 19.4, p = 1.32 x 10^-15^, Figure 2B). The distribution of allelic effects on the trait showed a huge effect on the mean PPV (mean equal contribution = 0.81, mean exponential = 0.32, F = 3409 p = 0, Figure 2C). The relation of mean proportion of detected TPL and number of migrants was approximately linear. The more migrants per generation, the higher the probability to detect more contributing loci. This effect was strong (F = 112, p = 2.35 x 10^-70^). The number of populations screened had a similarly strong effect on proportion of TPL among the selected loci (F = 117,4 p = 4.70 x 10^-95^). The values ranged from a mean PPV of 0.35 (s.d. = 0.21) with 12 populations to over 0.63 (s.d. = 0.32) with 60 populations. Given the chosen threshold, a diminishing return was observed above 36 populations sampled (Figure 2E). The phenotypic range in a simulation run had a low (r = 0.11, p = 3.02 x 10^-11^), yet significantly positive effect on detection of TPL. The realised range of population trait means in the simulations covered on average 16% (range = 0.1-57%) of the possible range. An increase of one unit in range increased the proportion of TPL by 0.02 (Figure 2F).

**Figure 2.**
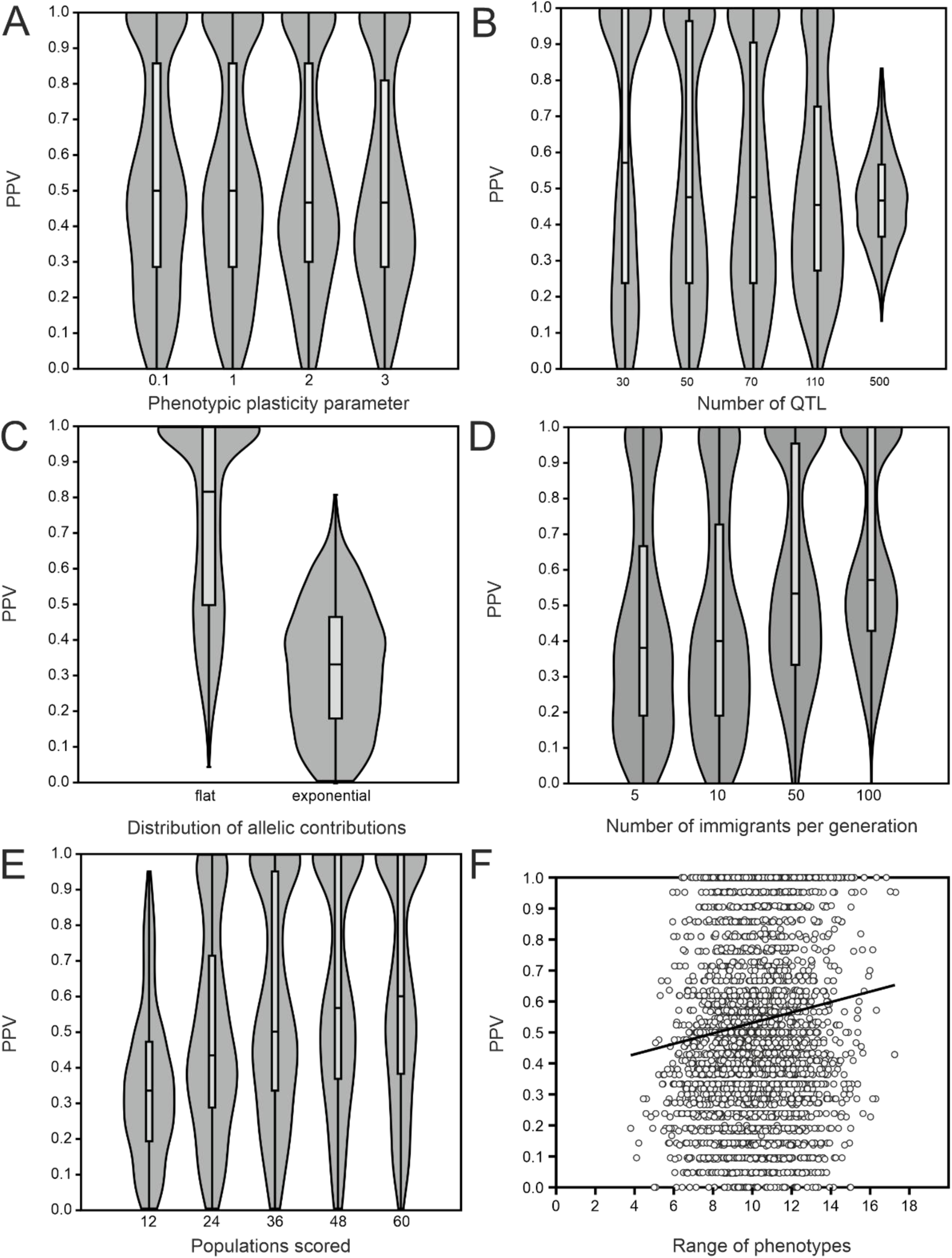
Effect of simulation parameters and emergent features on the proportion of identified true positive loci. A) Environmental variance parameter as a proxy for heritability. B) Number of QTL. C) Distribution of allelic contributions to phenotypic trait. D) Number of reproducing immigrants per generation. E) Number of populations scored for population phenotypic means and allele frequencies. F) Range of population phenotypic means as an emergent feature.

When jointly considering the effect of all parameters on PPV in a GLM, it turned out that all had a significant effect (Table 4). Their relative influence increased from phenotypic range (r² = 0.03) over heritability(r² = 0.06), number of QTL (r² = 0.07), F_ST_ (r² = 0.19), number of populations (r² = 0.20) to the number of allelic contributions, that had by far the greatest influence (r² = 0.45). In total, the parameters explained 66.4% of variance.

**Table 4.** Generalised Linear Model of factors influencing the proportion of TPL among outliers (PPV) in simulations.

| Factor | Coefficient | Std.err. | t | p | $r^2$ |
| --- | --- | --- | --- | --- | --- |
| Constant | 0.33 | 0.02 | 18.00 | <2e-16 |  |
| range_pheno | 0.01 | 0.00 | 4.02 | 0.00006 | 0.03 |
| heritability | -0.14 | 0.01 | -10.10 | <2e-16 | 0.06 |
| $n_{\text{qtl}}$ | 0.00 | 0.00 | -10.63 | <2e-16 | 0.07 |
| $F_{\text{ST}}$ | -5.72 | 0.19 | -29.84 | <2e-16 | 0.19 |
| $n_{\text{pop}}$ | 0.01 | 0.00 | 28.47 | <2e-16 | 0.20 |
| allelic_contr | 0.43 | 0.01 | 73.35 | <2e-16 | 0.45 |

### Hierarchical population structure

The introduction of a hierarchical population structure in simulations with a restricted parameter set did not change much with regard to the relative influence of the parameters (Supplemental Figure 6 A-F). The initial divergence parameter had no effect on PPV. The number of QTL loci (F = 12.13, p = 7.38 x 10^-6^), the distribution of allelic contribution (F = 888, p = 7.73 x 10^-109^) and the number migrants per generation (F = 18.7, p = 1.88 x 10^-5^) had a significant effect, just like the number of regions, going along with a growing number of subpopulations (F

= 4.06, p = 0.02). The final overall F_ST_ had a significant influence (p = 2.1 x 10^-5^), albeit with a relatively low effect size (r = -0.20).

### Minimum Entropy Feature Selection and statistical phenotype prediction

Minimum Entropy Feature Selection (MEFS) removed on average 8.72 (range = 2-14, s.d. = 4.14) loci, corresponding to a proportion of 0.38 (s.d. = 0.19) from the initally chosen outlier set. The procedure removed on average a larger proportion of FP than TPL (mean difference 0.14, t

= -19.9, p = 6.7 x 10^-79^). This increased the proportion of TPL in the final prediction set on average by 0.05 (range = -0.23-0.48, s.d. = 0.09) to a mean of 0.66 (range = 0-1, s.d. = 0.29, Figure 3).

**Figure 3.**
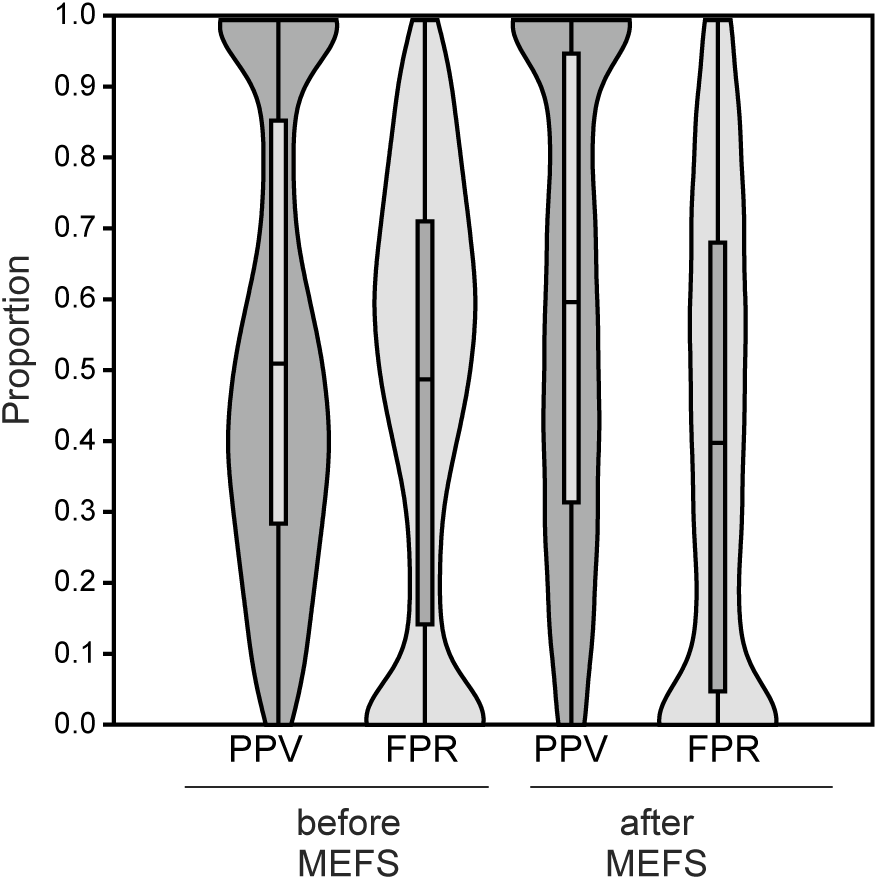
Effect of Minimum Entropy Feature Selection (MEFS) on the proportion of TPL and FP in the selected set.

The predictive accuracy of the SNP loci sets selected by MEFS was on average r = 0.61 (s.d. = 0.48). It ranged from -0.96 to 1.0. The distribution was highly skewed with 75% being higher than 0.37, the median was found at 0.86 and still 25% being higher than 0.97 (Supplemental Figure 4B).

The accuracy of mean population phenotype prediction depended linearly on the number of TPL in the prediction set (r² = 0.28, p = 0), with any additional TPL increasing the correlation coefficient by 0.05. Inversely, the accuracy of prediction decreased with a rising number of FP, but even with a considerable number of FP in the prediction set, accurate prediction was possible in a large number of cases. Overall, the prediction accuracy increased with increasing proportions of TPL among the prediction set, although even 100% TPL in the prediction set did not guarantee a highly accurate prediction (r > 0.8) in all cases.

As the prediction accuracy depended on the proportion of selected TPL, their relation to the individual simulation parameters was very similar to the results described in the previous section, except for the distribution of allelic contributions where the prediction accuracy was better for loci with exponential effects (Figure 4A-F). The number of populations screened was the most important factor. With 36 or more populations screened, 97.8% of simulations showed a prediction accuracy of 0.8 or better, independent of the other simulation parameters applied. In a GLM with all factors simultaneously considered, the proportion of TPL selected had the largest influence on prediction accuracy (r² = 0.28), followed by F_ST_ (0.19), the number of QTL (0.18), distribution of allelic contributions (0.17), the number of populations screened (0.11). Heritability had only a minor influence on the prediction accuracy (0.06), while the range of phenotypes was not relevant (Table 5).

**Figure 4.**
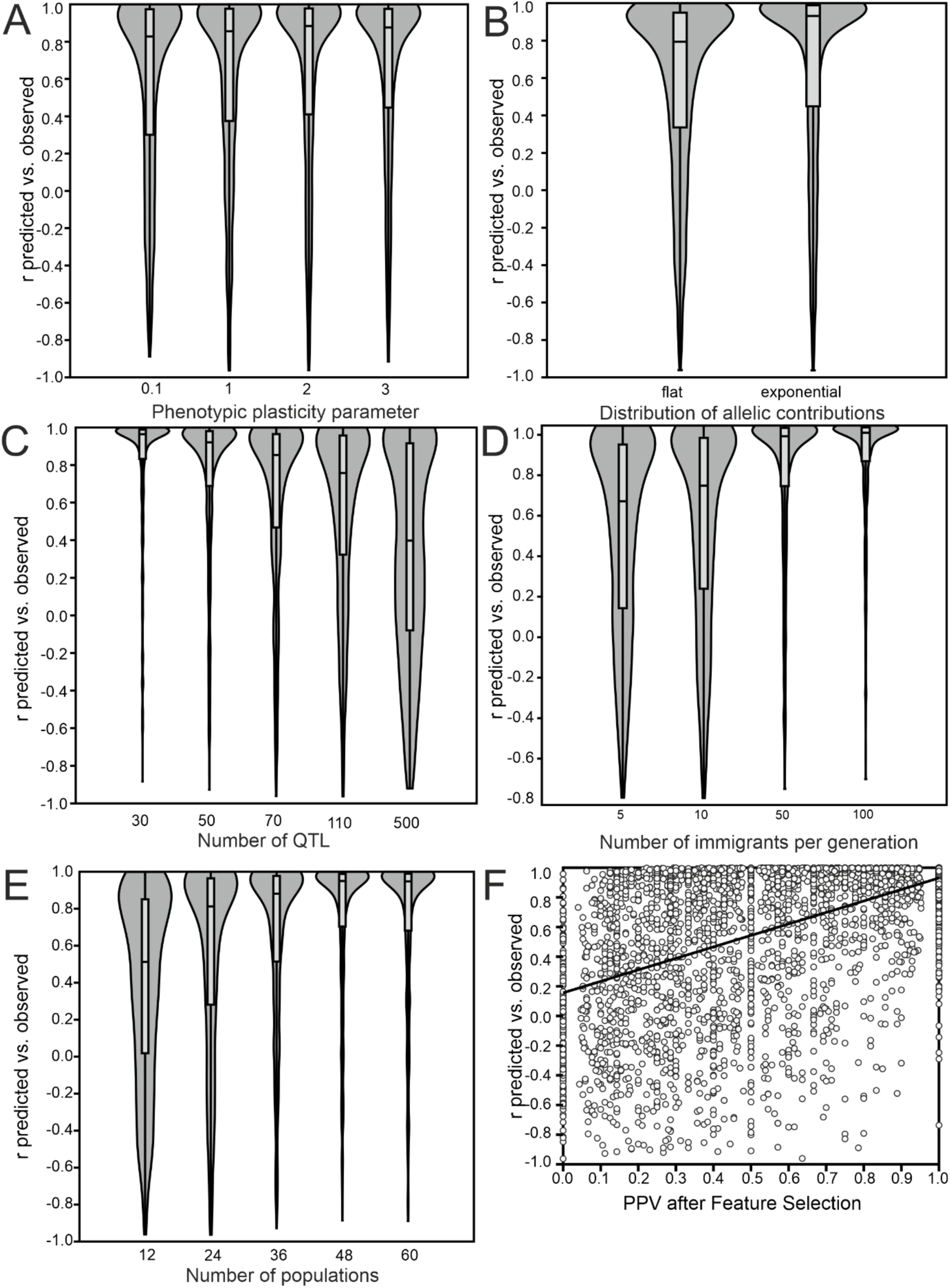
Influence of simulation parameters on the accuracy of statistical population mean phenotype prediction. A) Environmental variance parameter as proxy for heritability. B) Distribution of allelic trait contributions. C) Number of trait-underlying QTLs. D) Generation of independent evolution as proxy for population structure. E) Number of populations scored. F) Proportion of TPL in the prediction loci set after MEFS.

**Table 5.** Generalised Linear Model of factors influencing the accuracy of statistical phenotypic population mean prediction.

| Factor | Coefficient | Std.err. | *t* | *p* | $r^2$ |
| --- | --- | --- | --- | --- | --- |
| Constant | 0.69 | 0.04 | 17.31 | <2e-16 |  |
| range_pheno | 0.01 | 0.00 | 1.61 | 0.109 | 0.02 |
| heritability | -0.16 | 0.03 | -5.65 | 1.77E-08 | 0.06 |
| n_pop | 0.00 | 0.00 | 10.05 | <2e-16 | 0.11 |
| allelic_contr | -0.21 | 0.01 | -15.93 | <2e-16 | 0.17 |
| n_qtl | 0.00 | 0.00 | -18.62 | <2e-16 | 0.18 |
| $F_{ST}$ | -7.63 | 0.42 | -18.04 | <2e-16 | 0.19 |
| PPV after MEFS | 0.51 | 0.02 | 24.50 | <2e-16 | 0.28 |

### Performance comparison to individual GWAS

In every single simulation, iGWAS performed worse than popGWAS. The approach based on individuals found on average 7.42 TPLs (PPV 0.13), compared to 12.15 (PPV 0.53) with popGWAS (Suppl. Figure 4A). This difference in means of 4.73 loci was highly significant in a pairwise t-test (t >3000, p = 0). The results for hierarchically structured population were similar (mean difference 5.7 loci, paired t-test t = 19.62, p = 1.6 x 10^-62^). iGWAS did not suffer from p-value inflation either (mean = 1.15 s.d. = 0.18, (Suppl. Figure 4B)). In more than 70% of simulations, it was not possible to calculate a valid phenotpic prediction model for individuals with the identified loci. For the remaining cases, the mean predictive accuracy for the individual’s phenotype was 0.079 (Suppl. Figure 4C).

### Method performance with realistic effective genome sizes

The values for AUC-ROC ranged between 0.067 and 0.833, for AUC-PR between 0.013 and 0.730. There was an interaction between the effective genome size and number of populations scored. According to both AUC measures, the method performed best, when the number of populations scored was high and the genome small (Figure 5). An at least satisfactory (> 0.66 for AUC-ROC and > 0.53 for AUC-PR) overall performance was observed for 24 populations for the smallest genomes considered (1,000), for 36 populations up to 30,000 independent loci and for genome sizes up to 100,000 for 48 and 60. The similar values in both statistics and the plots suggested that there are diminishing returns for samples larger that about 48 populations. Moreover, closer inspection of the corresponding plots (Figure 6) suggested that for samples of 48 and 60 populations, an optimal ratio between TPL and FPL exists for approximately the 25 highest outlier loci, independent of genome size. For combinations with good performance, the maximum F1 score suggested that choosing the 30 highest outlier provided the optimal compromise between maximising TPR and minimising FPR (Supplemental Figure 7).

**Figure 5.**
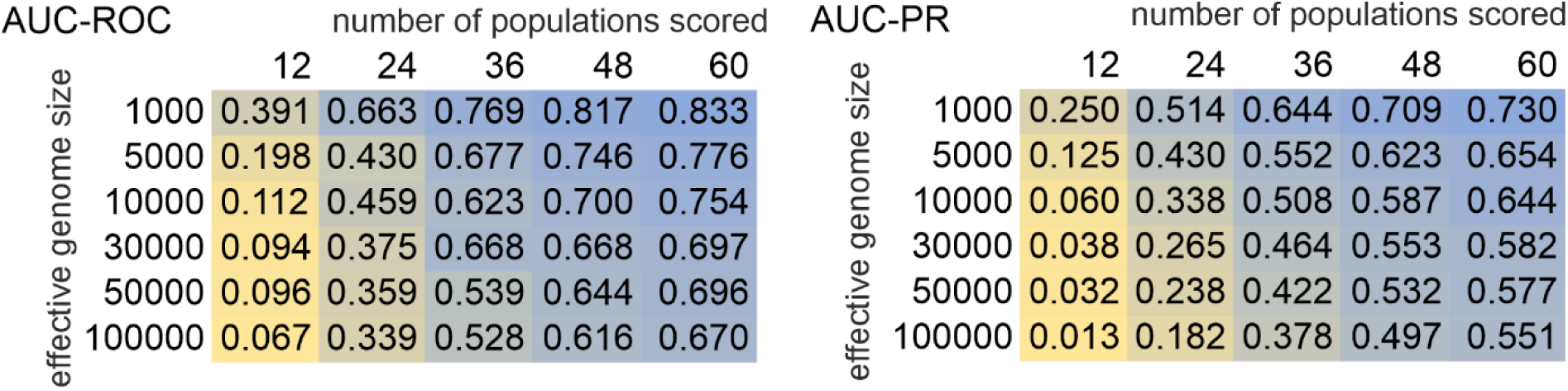
Heat-map of AUC-ROC (area under the curve – receiver operator characteristics) and AUC-PR (area under the curve – precision recall) in relation to effective genome size and number of populations scored.

**Figure 6.**
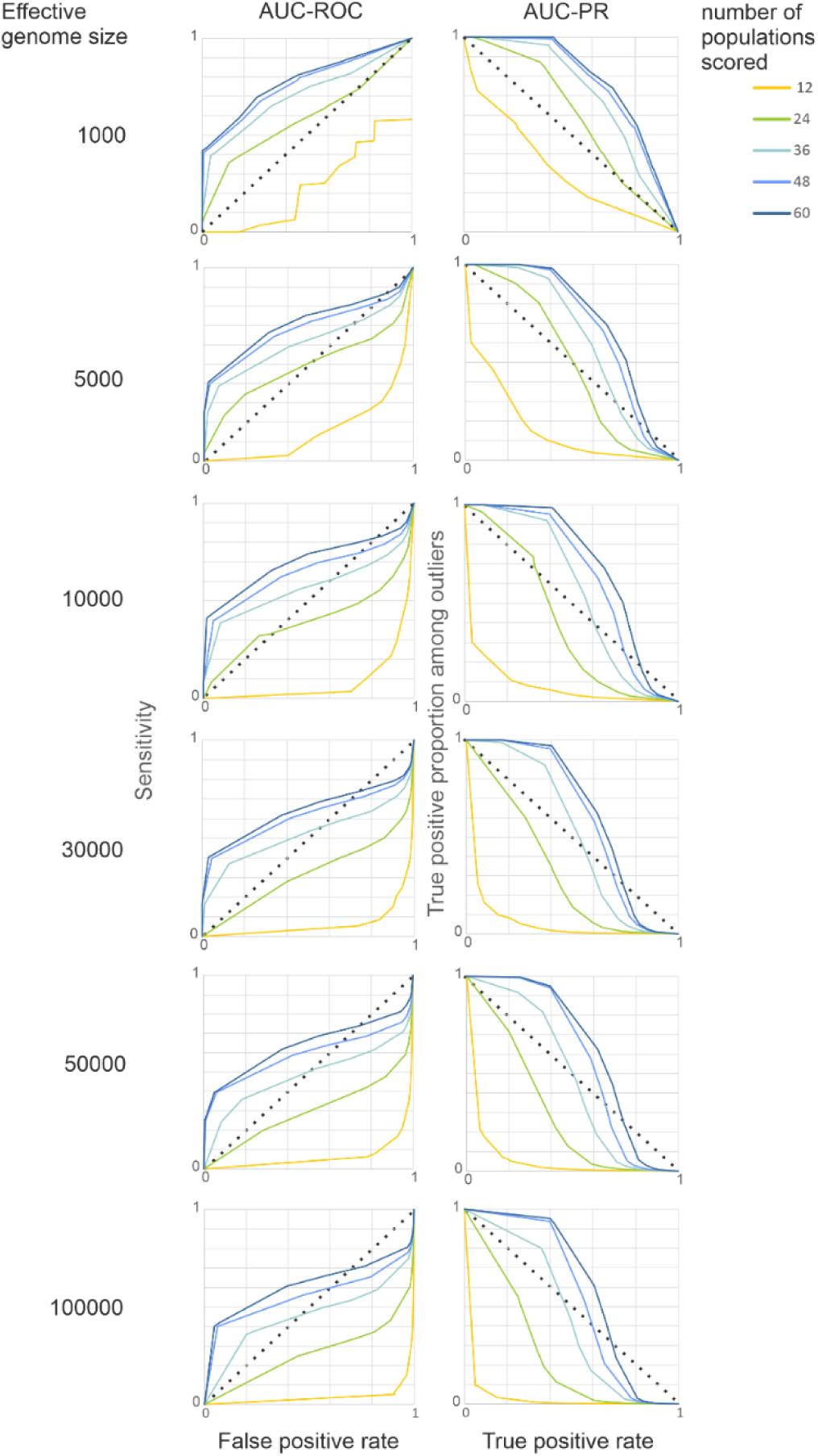
AUC-ROC and AUC-PR for a range of effective genome sizes. In the left column are the plots of AUC-ROC, i.e. FDR on the x-axis versus TPR on the y-axis. The right column shows AUC-PR plots, i.e. TPR on the x-axis versus PPV on the y-axis. The dotted lines indicate the threshold for a random effect.

## Discussion

This study used extensive forward simulations to explore the potential of a novel GWAS approach utilising phenotypic population means and genome-wide allele-frequency data to identify loci potentially underlying the population differences in quantitative polygenic traits and predict the mean phenotypes of unmeasured populations. While the approach seems to be generally useful in a wide range of cases, there are also clear limits to its applicability.

### General validity of the underlying assumptions

The first major goal of the study was understanding the conditions under which the hypothesised pattern of a linear relation between the population allele frequencies at causal loci and the phenotypic population means of the respective trait emerges. The initial simulations demonstrated that the expectation for the covariance of random population “allele frequencies” at contributing quantitative trait loci (QTL) and the respective population trait mean is consistently positive when an additive model applies. This is an inherent consequence of the common dependence of both variables on the QTL genotypes of the individuals in a population, as demonstrated in (3). Additive models seem to be an appropriate statistical approximation for most quantitative traits at population level (Hill et al., 2008), despite the description of many epistatic interactions on the molecular level (Moore & Williams, 2005).

The relation appeared to be largely independent of the distribution shape of locus contributions to the trait. While in the case of equal contributions i.e. a flat distribution, the correlation coefficients of individual loci are themselves a random variate, normally distributed around the expected mean. As the distribution becomes increasingly skewed, locus contribution becomes predictive of the correlation to the trait. Loci contributing more to the trait and thus accounting for more of the phenotypic variance will likely have a higher correlation of their allele frequencies to the population mean. Conversely, the expectation for non-contributing loci is approaching zero. Therefore, it is principally possible to exploit the correlation between allele frequencies and population trait means for the identification of loci underlying an additive quantitative trait.

However, some statistical limitations became obvious. Firstly, as the number of QTL increases, the expected mean correlation coefficients become so small that they are likely to be indistinguishable from the tail of the zero-centered normal distribution of non-contributing loci, even with an unrealistically high number of samples. Consequently, the method for identifying QTL by the positive covariance of their allele frequencies with the population trait means is *a priori* more suited for oligogenic to moderately polygenic traits. Secondly, the number of QTL and the distribution of locus contributions may influence the statistical identifiability of individual QTL. In particular, loci that contribute only minimally to the trait or that fall by chance below the expected mean correlation coefficient may overlap with the tail of the distribution of non-contributing loci.

These predictions remind of similar conditions for the contribution of different QTL architectures to phenotypic adaptation described by (Höllinger et al., 2023). They assert that phenotypic adaptation of oligogenic traits is achieved by detectable allele frequency shifts at some but not very many loci, while adaptation in highly polygenic traits is rather achieved by subtle perturbations of standing variation, with respective consequences for their detectability. Just as expected here, they stress the importance of stochastic effects that may lead to apparently heterogeneous locus contributions (Höllinger et al., 2023).

Interestingly, deviation from a selection-drift-migration equilibrium were not an impediment for detecting contributing loci. Best possible adaptation was regularly achieved within less than 10 generations. True to theoretical expectations, higher gene-flow among populations led to less perfect local adaptation for a given selective strength (Brady et al., 2019). Drift-migration balance was not reached within a reasonable number of generations. This was, however, not relevant for popGWAS performance, which was mainly driven by the degree of population differentiation, everything else being equal. This insensitivity to equilibrium conditions is an advantage of the method as natural populations tend not to be in equilibrium (Müller et al., 2022). Rather, natural populations are constantly tracking variable selective optima (Pfenninger & Foucault, 2022; Rudman et al., 2021), which leads to intermediate allele frequencies at the loci under such a selective regime (Höllinger et al., 2023). However, performance of the method under particular demographic scenarios like extreme bottlenecks or sudden range expansions remain to be tested.

### Limiting factors for the identification of contributing loci in natural settings

The Wright-Fisher forward simulations of a quantitative trait in a subdivided population with realistic properties and sample sizes largely confirmed the theoretical expectations. In particular when a sufficient number of populations was scored (in this case more than 36), a large proportion of true positive loci could be reliably identified, with the exception of a few parameter combinations. The genetic architecture of the trait was an important predictor for the ability to identify causal loci. While loci underlying oligogenic and moderately polygenic traits could be fairly reliably identified, the highly polygenic scenario tested (500 loci) performed poorly. A higher proportion of TPL was identified when the locus contribution to the trait was identical than in the case of an exponential allele effect distribution This was likely due to the tendency of higher correlations between large effect loci and the trait, which allowed the loci from the tail of the distribution to vary freely, making them indistinguishable from non-contributing loci.

The influence of mean heritability on GWAS performance was not marked. Even down to trait heritability estimates of 0.3, the success rate was only slightly reduced. This effect may be attributed to the averaging of phenotypes and genotypes across multiple individuals, which is likely to mitigate the inherent noise associated with individual data (Johri et al., 2022; Stinchcombe & Hoekstra, 2008). This finding is consistent with observations by (Zhang et al., 2018), who employed pooled data for GWAS. From a practical standpoint, the findings suggest that inevitable errors in phenotyping, which can compromise GWAS performance on individuals (Barendse, 2011), are likely to be less problematic when using the mean measured over many individuals. Furthermore, this finding indicates that the failure to entirely remove non-additive variance from the analysis does not necessarily compromise the method’s ability to reliably identify trait-associated loci.

From a statistical perspective, it was anticipated that the range of phenotypic population means would influence the identification of true positive loci to some extent, given that a larger range of phenotypic means is inherently associated with on average larger allele-frequency differences among populations. The choice of populations with a large range of phenotypic variance is therefore crucial. It is, however, important to emphasise that the underlying causes of the observed differences in trait means among populations are not of primary concern. These may be attributed to local adaptation, but also to maladaptation, human choice, or other factors. Likewise, increasing the number of populations screened increased the statistical power of the approach. However, it seemed that increasing the number of samples led beyond a certain threshold to diminishing returns in statistical power gain.

### Effect of population structure on the detection of contributing loci

It is long-since known that population structure can have a confounding effect on GWAS studies (Marchini et al., 2004). Confounding associations between bi-allelic SNPs and a phenotype may easily arise in iGWAS, because phenotypically more similar individuals may be also genetically more similar because they share common ancestry as members of the same family/population/clade. In addition, a quantitative or qualitative phenotype (y) in iGWAS is related to a genotype (x) either by a linear or logistic model – with exactly three possible values of x. This leads inherently to low statistical power, requiring a very high number of individuals (O’Connor, 2021) and extensive corrections for population structure to obtain a reliable statistical association (Sul et al., 2018). popGWAS follows a principally different approach; the possible values of x are allele frequencies in different populations and thus limited only by sampling effort. This gives the approach a strong inherent statistical advantage, which might be also the reason that the method does not suffer from p-value inflation.

Population structure can nevertheless interfere with reliable associations in popGWAS, albeit for different reasons. A pronounced population structure (F_ST_ > ∼ 0.07) was indeed a major factor impeding reliable identification of true positive loci, even with a high number of samples and regardless of the reasons for the structure. However, this was probably more due to distinct evolutionary trajectories in independently evolving populations. If the evolutionary trajectory of the populations analysed is sufficiently independent, either by drift and/or differential availability of adaptive mutations, similar phenotypes may have a differential genomic basis, allowing at best to identify any common variants. The genetic redundancy of polygenic traits can lead to evolution of the same phenotypes from different genomic bases (de Vladar & Barton, 2014; Kaneko & Furusawa, 2006), even if evolving from the same ancestral population (Barghi et al., 2019, 2020; Pfenninger et al., 2015). If different loci in different populations are causal for the observed phenotypic differences, a linear relation between population means and allele frequencies is not to be expected. It is therefore important that the allele frequencies in the studied populations are correlated either by recent common descent and/or recurrent gene-flow, i.e. that the population structure between the population scored is weak (Mathieson, 2021).

By relating allele frequencies with mean phenotypes false associations may also arise, if the trait under scrutiny is effectively neutral and thus the constituting loci follow the overall drift pattern or the ecological differences causing the phenotypic differences are associated to an isolation-by-distance pattern. In this case, the matrix of mean phenotypic population differences (or Q_ST_) should show a strong positive correlation with the overall F_ST_ matrix among populations. A trait showing this easily testable pattern may therefore *a priori* not be suitable for analysis with the method. To avoid such a situation, it is recommended to test for (the absence of) a correlation between genome-wide genetic distance and differences in phenotypic means (e.g. by a Mantel’s test).

### Accurate statistical genomic prediction in a wide range of conditions

Genomic prediction is deemed to be one of the major tools for the mitigation of climate change on biodiversity (Aguirre-Liguori et al., 2021; Bernatchez et al., 2023; Capblancq et al., 2020; Waldvogel et al., 2020). One of the major findings of the presented approach was the highly accurate prediction of the phenotypic population means from genomic data above a certain number of populations screened. Given the limited performance of genomic prediction in a medical context (Visscher et al., 2017) this was initally quite surprising. However, it should be *a priori* easier to predict the differences in population means compared to differences among individuals for the following reasons. The difference in means among subpopulations was mostly governed by only a couple of loci, because the range of phenotypic population means extended over only a relatively small part of the theoretically possible range. This applies to the simulations but most likely also in nature it is improbable to encounter populations that are (nearly) fixed for the respectively alternative alleles over all or even most trait contributing loci and show an accordingly extended phenotypic range. Consequently, already the allele frequency differences of a couple of loci actually need to follow a linear model to explain most of the variance among phenotypic population means – the rest is free to vary and behave (almost) like neutral loci. Even the fit of the highest outlier loci to a linear model was far from perfect as was expected from the genetic redundancy of a polygenic trait architecture (Barghi et al., 2020). However, as long as these important loci are the same over the majority of subpopulations (either due to common descent or gene-flow), accurate predictions of population trait means from these loci is therefore possible. Such loci likely have an intermediate frequency, because alleles of very low or very high frequency (i.e. present only in few or almost all individuals with not necessarily extreme phenotypes) by definition cannot have a large impact on the phenotypic population mean. This is true even when the rare allele is of large effect for a respective allele-carrying individual. Therefore, a locus whose AF is important for the difference in means among populations might have rather low predictive power for the individual, because as a locus of likely intermediate frequency, all genotypes are necessarily realised with high probability within a population. The accurate prediction of individual phenotypes thus likely requires much more loci than the prediction of the phenotypic population mean.

Finally, it is crucial to distinguish between two major goals of GWAS: i) find the loci functionally underlying biologically relevant variation and ii) the accurate prediction of untyped instances. As (Shmueli, 2010) pointed out, the goals of explaining a phenomenon on the one hand and predicting it on the other are related but not identical. In particular it is important to note that an accurate prediction model does not need to contain all features that are necessary for a comprehensive functional description of the system (Shmueli, 2010). For accurate prediction, it is not even necessary that all features in the prediction model have actually a functional role, as long as they are statistically strongly associated (Shmueli, 2010). While the results have shown that prediction tends to become more accurate if more truly functional loci are included, it is therefore not necessary that the prediction model includes all or even most of the functional loci.

Contrary to its application in medicine or selective breeding (Wray et al., 2019), however, accurate prediction of population responses is in many instance of the current biodivsity crisis probably more important than the prediction of individual phenotypes. Within the limits outlined above, the proposed method delivered in many instances very accurate predictions (r > 0.8 in > 90% of cases for more than 36 populations) of population mean phenotypes. It should be noted, however, that the prediction is statistical in the sense that it produces a prediction score (de Los Campos et al., 2018) that correlates with the mean population phenotype and is explictely not a comprehensive functional quantitative genetic model of the trait Just like with any other genomic prediction (Kachuri et al., 2024), this limits the transferability of the prediction to other, more distantly related lineages or species.

Reducing the false positive rate before prediction is in any case advisable, as it proved to be the most important factor of prediction success with independent data. The application of a Machine learning approach, in this case Minimum Entropy Feature Selection (MEFS), prior to prediction effectively reduced the already low false positive rate among the initially selected loci further. Whether other approaches would yield even better results remains to be tested. Other population genetic parameters influenced prediction success in a very similar fashion as the initial true positive rate. One notable exception was distribution of locus contributions. While true positive loci were more reliably identified from a flat distribution, prediction worked better when many loci of large effect were among the prediction set, most likely because these loci contribute more to phenotypic variance among populations (Jain & Stephan, 2015).

### Typical genome sizes of real species are no obstacle

The perhaps most important challenge was showing that the proposed method has enough statistical power to distinguish at least a part of the unknown, but likely relatively small number of QTL reliably from the large number of non-contributing loci in real genomes of real species. The evaluation of method performance with AUC-ROC and AUC-PR, as recommended recently (Lotterhos et al., 2022), showed a satisfactory performance even for genomes with moderately high effective sizes, provided a sufficiently high number of populations is screened. In particular restricting the selection of potentially causal QTL on a few dozen of the highest outliers promises to yield very low false positive rates. However, as the success rate depended very much on the biological characteristics like genome size, population structure, LD and so on, of the investigated taxon, no general recommendations can be given here, except to carefully take them into account. As shown above, already a limited number of true positive loci may be sufficient for reliable genomic prediction.

### Practical considerations

The proposed method finds rather genomic regions or haplotypes associated to the trait in question than directly causal SNPs. However, this is true for most GWAS methods (Wang et al., 2010) and therefore fine-mapping and inference of causal processes remain to be done (Wallace, 2021). In practice, this requires that regions with high SNP outlier density need to be collapsed to haplotypes prior to further analysis. Knowledge on the local LD-structure, mean haplotype length, respectively recombination landscape can aid haplotype identification (Flister et al., 2013). Recently developed machine learning approaches makes such information available for pooled data (Adrion et al., 2020).

The possibly largest advantage of the proposed method is therefore its data efficiency, if pooled sequencing is applied. Because the Pool-Seq approach (Schlötterer et al., 2014) yields highly accurate estimates of genome-wide allele frequencies at SNP sites (Czech, Peng, Spence, Lang, Bellagio, Hildebrandt, Fritschi, Schwab, Rowan, & consortium, 2022) the necessary sequencing effort is marginal compared to individual based approaches (Ziyatdinov et al., 2021). This is mainly due to the relatively reduced costs for library preparations necessary that, depending on the genome size, often make up a large share of or even exceed the actual sequencing costs in individual analysis. Moreover, the popGWAS method out-performed iGWAS in all cases, if the same sequencing effort is applied as a benchmark.

The proposed method makes GWAS studies accessible for the usual funding in the field of biodiversity. Pooled sequencing for GWAS has been proposed (Yang et al., 2015) and applied (Giorello et al., 2023; Kumar et al., 2022; Pfenninger et al., 2021) with extreme phenotypes. Automated large scale phenotyping is a very active field of research, with satellite or other remote sensing approaches, flow cytometry, bulk measurements of metabolic rates, metabolomes, transcriptomes, pooled HPLC or mass spectrometry, automated video-based phenotyping, mass CT scanning provides already access to large amounts of data. Sampling the required number of demes or subpopulations can pose a challenge for certain taxa, however, e.g. biodiversity monitoring schemes or museum collections may offer readily access to respective samples. Application of the method to a real world data set from European beech on phenological traits (Pfenninger et al. in prep), showed that such an approach can be sucessfully performed with moderate logistic effort. Many of the identified loci by popGWAS were previously implicated in phenological traits and yielded excellent predictive power in predicting the mean phenotypes of independent populations (Pfenninger et al. in prep).

## Conclusion

This study demonstrated the potential of the proposed GWAS approach for biodiversity genomics. As with any GWAS method, its usefulness depends on the study goals. If it is the aim to infer all loci in a genome contributing to a given quantitative trait and to quantify their relative influence, probably no existing (single) GWAS method and no experimental design can achieve this. If it is the goal to accurately predict individual quantitative phenotypes with the inferred genomic basis, e.g. for medical or breeding purposes, the suggested method may not lead very far, as detailed above. Nevertheless, it can add a substantial number of candidate loci that are not easily found by other methods. However, if it is important to infer the major contributing loci for quantitative differences in ecologically relevant traits observed among natural populations, popGWAS can be a valuable tool. By carefully considering the factors influencing its performance and addressing the limitations, this method holds some promise in identifying the genetic basis of complex traits and performing accurate statistical predictions of phenotypic population means from genomic data in natural populations.

## Acknowledgements

The author wants to thank Bob O’Hara, Barbara Feldmeyer and the reviewers for helpful comments on the manuscript.

## Funding

The author declares that he has received no specific funding for this study.

## Conflict of interest disclosure

The author declares that he complies with the PCI rule of having no financial conflicts of interest in relation to the content of the article.

## Data, scripts, code, and supplementary information availability

Supplementary information including the Python code used for the simulations is available at https://10.5281/zenodo.15006644

## Notes

https://10.5281/zenodo.15006644

### Competing Interest Statement

The authors have declared no competing interest.

### Summary of Updates

This is the final version as peer-reviewed and recommended by PCI EvolBiol

